# A microvascularized *in vitro* liver model for disease modeling and drug discovery

**DOI:** 10.1101/2024.07.05.602196

**Authors:** Flavio Bonanini, Roelof Dinkelberg, Manuel Caro Torregrosa, Nienke Kortekaas, Tessa M. S. Hagens, Stéphane Treillard, Dorota Kurek, Vincent van Duinen, Paul Vulto, Kristin Bircsak

**Author notes:** Contributed equally (Flavio Bonanini, Roelof Dinkelberg).

## Abstract

Drug discovery for complex liver diseases faces alarming attrition rates. The lack of non-clinical models that recapitulate key aspects of liver (patho)-physiology is likely contributing to the inefficiency of developing effective treatments. Of particular notice is the common omission of an organized microvascular component despite its importance in maintaining liver function and its involvement in the development of several pathologies. Increasing the complexity of *in vitro* models is usually associated with a lack of scalability and robustness which hinders their implementation in drug development pipelines. Here, we describe a comprehensive liver MPS model comprising stellates, liver-derived endothelial cells and hepatocytes conceived within a scalable and automated platform. We show that endothelial cells self-organize in a microvascular network when co-cultured with stellates in a hydrogel. In a tri-culture, hepatocytes polarize accordingly, with a basolateral side facing blood vessels and an apical side facing bile-canaliculi-like structures. Stellates interact and surround the hollow microvessels. Steatosis was induced by exogenous administration of fatty acids which could be prevented by co-administration of firsocostat. Administration of TGF-β resulted in an activated stellate cells phenotype which could be prevented by the co-administration of SB-431542. The model was implemented on a microtiter plate format comprising 64 chips which enabled the development of a fully automated, multiplexed fibrosis assay with a robust Z’ factor suitable for high-throughput applications.

## Introduction

Developing therapies for complex liver diseases is a prime goal of today’s pharmaceutical industry. Yet, attrition rates are high as therapies for complex diseases such as metabolic dysfunction-associated steatohepatitis (MASH) are historically difficult to develop. A root cause underlying these attrition rates is the lack of models capturing human pathophysiology in sufficient detail. In fact, conventional non-clinical models predominantly comprise animals and *in vitro* cell monocultures (1,2). Whereas the first suffers from interspecies translational limitations, the latter lacks the complexity of human physiology. In many cases homeostasis and diseases occur in the interplay between hepatocytes, non-parenchymal cells (NPCs), such as stellate cells, endothelial cells and resident macrophages (3–8). Hence, researchers have developed various advanced *in vitro* liver models following a more comprehensive approach where hepatocytes and several NPCs are combined. Multicellular spheroids, for example, offer a great degree of robustness and scalability and can show desirable functionality (9–17). Adult and fetal stem cell derived organoid approaches have shown expandability of donor-derived epithelial cells which promise translational potential (18–20). Yet, these organoids are comprised of exclusively epithelial cells and interfacing them with NPCs remains largely unexplored. Finally, iPSC derived liver organoids have been demonstrated to mimic embryonic development, differentiating into a large subset of NPCs in addition to parenchymal cells (21–23).

Although proven useful for several applications, each of these approaches have several limitations, such as disorganized cellular arrangements, lack of NPCs, require cumbersome and lengthy protocols or show limited maturation. Notably, all these systems lack a functional and organized vascular network and are hence limited in reflecting the spatial organization of the liver microstructure.

Microphysiological systems (MPS) provide a better physiological context by incorporating a three-dimensional (3D) environment, extracellular matrix, fluid flow and the appropriate cellular composition and organization found *in vivo* in microfluidic chips (24–27). MPS offer therefore the possibility of generating accessible, vascular systems which could potentially be used to achieve liver sinusoid-like structures, with stratified cellular arrangement as found within the native organ. This has been attempted following several approaches. One approach consists of generating a hollow channel within a hydrogel that can be perfused and populated with endothelial cells forming a perfusable blood vessel. Hepatocytes can be interfaced to the artificial blood vessel and this has been shown to improve hepatocyte function, as well as increasing sensitivity for drug toxicity testing (28). Perfusable liver sinusoids have also been developed by microneedle-assisted fabrication enabling a dual blood supply, resulting in increased cell viability and hepatocyte metabolism (29). The authors also report the spontaneous emergence of secondary sinusoids, implying that in the right context, cells possess considerable degree of self-organization capacity. Others have attempted to recreate the stratified cellular organization by confining different cell types in different microfluidic compartments, such as sinusoidal endothelial and Kupffer cells with a porous membrane separating stellate cells and hepatocytes (30). This resulted in increased hepatic function, as well as allowing neutrophil recruitment upon exposure to inflammation stimuli.

Yet, the implementation of liver MPS in drug development remains limited. The primary challenges include operational complexity, lack of compatibility with automation processes, manufacturing and material limitations, and lack of a standardized format.

We previously described a workflow for the vascularization of hepatic spheroids and organoids (31). The concept utilizes a commercially available microfluidic platform consisting of 64 microfluidic chips patterned underneath a 384-well microtiter. We demonstrated the impact of liver co-culture on the remodeling of the vascular bed and were able to mimic liver veno-occlusive disease. Yet, the paper did not report a full equivalent of a liver sinusoid, lacking hepatic stellate cells, liver sinusoidal endothelium and polarized hepatocytes.

Here, we report a liver MPS model with sinusoid-like structures for drug discovery and development. We demonstrate that primary liver endothelial cells can form microvascular network when embedded within a fibrin matrix. Inclusion of induced pluripotent stem cell-derived hepatocytes (iHEPs) alongside primary liver endothelial cells and stellate cells results in the formation of sinusoidal-like structures, with direct contact between hepatocytes and NPCs. Hepatocytes self-assembled into plate-like structures with bile-blood polarization. To demonstrate the applications of our liver MPS, we developed a steatosis assay by inducing lipid accumulation in the hepatocytes by exogenous administration of free-fatty acids and showed that these could be ameliorated by the addition of Firsocostat. We set up an automated TGF-β-induced fibrosis assay with multiplexed readouts for liver function, viability, and vascular network integrity. The assay shows an excellent window for drug screening applications. To our knowledge, the structural relevance of this model in combination with its ease of use, throughput and scalability has not yet been shown in other works and poses a fundamental advancement of liver MPS models for drug development.

## Methods

### Subculture of primary LDECs and HSCs

T-75 cell culture flasks (Corning #734-2705) were used for subculturing of primary hepatic stellate cells (HSCs) (ScienCell #5300) by pre-coating with 150 µL 0.1mg/mL Poly-L-Lysine + 4850uL PBS (R&D Systems #3438-100-01). T-75 flasks used for subculturing primary liver-derived endothelial cells (LDECs) (ScienCell #5000) were pre-coated with 150 μL 1 mg/mL Fibronectin + 4850 μL PBS (Sigma Aldrich #F1141-5MG) 2 hours prior to cell thawing. LDECs and HSCs were thawed from cryostorage on a waterbath at 37°C and directly seeded into the flasks. HSCs and LDECs were cultured in complete SteCM (ScienCell #5301) and ECM (ScienCell #1001) media respectively for 5 days with a full medium change every 2-3 days in a humidified incubator (37°C 5% CO_2_). Cells were then dissociated after a pre-wash with HBSS (Thermo Fisher #14175095) and incubated for 3-5 minutes at 37°C with a 1:10 dilution of trypsin 2.5% (Thermo Fisher #15090-046) in PBS (HSCs) or undiluted trypsin 2.5% (LDECs). Trypsin was neutralized with 5 mL of Trypsin Neutralizing Solution (TNS, Lonza #CC-5002). Content was collected into separated 50 mL tube (brand) containing 10 mL of fetal bovine serum (FBS; Gibco # 16140-071). Additional 5 mL of TNS was added to each flask to rinse and collect any remaining cells and cells were counted using the trypan blue exclusion method. Cell solutions were then centrifuged at 200g for 5 minutes after which they were resuspended in cold freezing medium composed of 70% culture medium, 20% FBS and 10% DMSO at 500’000 cells/mL. Cells were cryopreserved at −150°C.

Prior seeding in the OrganoPlate, one cryovial was thawed for each cell type and cultured as described above for 5 days. After dissociation, cells were centrifuged and resuspended in 1 mL of complete Dulbecco’s Modified Eagle Medium (DMEM, Gibco #11965-092) containing 10% FBS, 1% penicillin-streptomycin (P/S, Sigma-Aldrich #P4333-100ML) and 1 mM sodium pyruvate (Thermo Fisher #11360070), here called DMEM+. Counting was performed using the trypan blue exclusion method. After cell counting, cells were centrifuged again for a second resuspension in desired volume of DMEM+.

### Subculture of non-parenchymal cells

Liver NPCs (Lonza #HUCNP, lot. No. NPC183061) were cultured in T-75 cell culture flasks coated with 50 µg/mL Purecol (Advanced BioMetrix, #5005-B). Cells were maintained in a humidified incubator (37°C 5% CO_2_) with a culture medium consisting of a 1:1 mixture of EMEM (ATCC #30-2003) + 10% FBS and Endothelial Cell Growth Medium 2 (PromoCell #C-22011). Medium was changed every 2-3 days and after reaching confluency, cells were dissociated using 3 mL TrypLE™ Express Enzyme (Thermo #12604021). Cells were counted using the trypan blue exclusion method, centrifuged 300g for 5 minutes and resuspended in desired volume of DMEM+.

### Subculture of primary human umbilical-vein endothelial cells

Primary human umbilical-vein endothelial cells (HUVECs, Lonza #2519AS) were cultured for a maximum of 4 passages cultured in EGM™-2 (Lonza #CC-4176) in T-175 or T-75 culture flasks (Thermo, #159910) in a humidified incubator (37°C 5% CO_2_). Cells were dissociated using TrypLE™ Express Enzyme at 37°C and cryopreserved at passage 4 in complete culture medium + 10% DMSO at 1’000’000 cells/mL. Passage 4 HUVECs were thawed and cultured for 5 days prior to seeding in the OrganoPlate.

### OrganoPlate Seeding

On the day of seeding, HSCs, LDECs or NPCs were collected as described in the subculturing sections. Fujifilm’s iPSC-derived hepatocytes ICell® Hepatocytes 2.0 (iHEPs, Cellular Dynamics C1090) were thawed following manufacturer’s instructions. Cells were mixed at appropriate densities with 5 mg/mL end concentration Fibrinogen (Enzyme Research Laboratories #FIB1) and 0.1 U/mL end concentration thrombin (Enzyme Research Laboratories # HT 1002a). The mixture was vigorously mixed and pipetted from a tube onto the OrganoPlate Graft by dispensing 1.35 μL through the graft chamber hole (B2 in Figure 1B) using a Sartorius P10 Pipette at minimum dispensing speed. The plate was then placed in a humidified incubator (37°C, 5% CO_2_) for 15 minutes. 50 μL of DMEM+ was added to the graft chamber. After 1-2 hours the medium in the graft chamber was replaced with iHEP plating medium prepared as described in ICell® Hepatocytes 2.0 culture protocol (here called iHEP medium) supplemented with 100 kIU Aprotinin (Nordic Group #16ND4110). HUVECs were then dissociated as mentioned in the subculturing protocol and resuspended at a density of 10000 cells/μL in ECGM-2 (Promocell #C-22011) supplemented with 100 kIU Aprotinin. 1 μL of this cell suspension was dispensed into the perfusion channels of each chip (A1 and A3 in Figure 1B). 50 μL of ECGM-2 + Aprotinin was added to the inlet wells (A1 and A3 in Figure 1B). The plate was then placed static in a humidified incubator (37°C 5% CO_2_) to allow endothelial cell attachment. After 2 hours, an additional 50 μL of ECGM-2 + Aprotinin was added to each perfusion channel outlet (B1 and B3 in Figure 1B) and the plate was placed on an OrganoFlow rocker (Mimetas B.V. #MI-OFPR-L) set at 14 degrees with 8-minute intervals to induce gravity-driven perfusion through the perfusion channels. Automated cell seeding followed an identical protocol but was performed using a Biomek i5 (Beckman Coulter).

**Figure 1:**
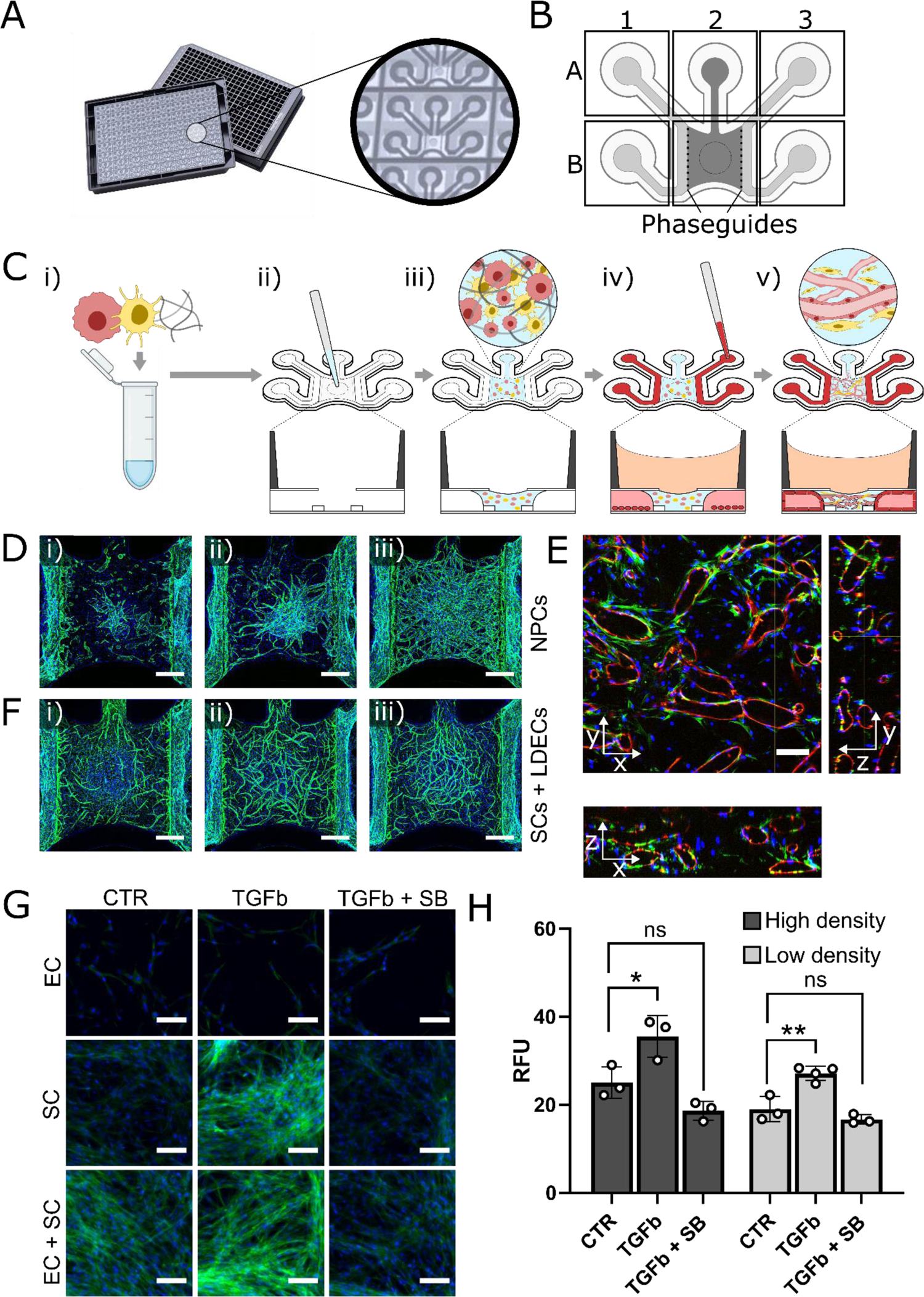
Establishment of a liver microvascular system. **(A)** Rendering of the OrganoPlate Graft, a microfluidic platform with 64 individual chips patterned underneath a 384 well-plate. **(B)** Schematic of a single chip, with Phaseguides separating two lateral channels with a central chamber. **(C)** Overview of chip seeding procedure. A mixture of cells and fibrinogen is prepared off chip (i) and pipetted into the opening of the Graft chamber (ii). The hydrogel is then allowed to polymerize (iii). Subsequently, HUVECs are introduced in the lateral channels (iv) and the culture is maintained on a rocking platform, allowing formation of lateral tubules and microvasculature within the hydrogel (v). **(D)** Confocal maximum intensity projections with VE-Cadherin staining (green) and nuclei (blue) showing microvasculature morphology after 10 days of culture of NPCs with seeding densities of 5416 (i), 8125 (ii) and 12188 (iii) cells / µL. Scalebar = 500 µm. **(E)** Confocal orthogonal views of a chip cultured with 12188 NPCs for 10 days stained for CD-31 (red), alpha-smooth muscle actin (green) and nuclei (blue). Scalebar = 100 µm. **(F)** Confocal maximum intensity projections with VE-Cadherin staining (green) and nuclei (blue) showing microvasculature morphology after 10 days of coculture of LDEC:HSC with seeding densities of 6500:325 (i), 6500:1625 (ii) and 6500:3250 (iii) cells / µL. Scalebar = 500 µm. **(G)** Confocal sum projections showing F-actin (green) and nuclei (blue) of HSC monoculture, LDEC monoculture and LDEC:HSC coculture after 24 hours incubation with 10 ng/mL TGF-b with or without 10 µM SB431542. Scalebar = 100 µm. **(H)** Quantification of F-actin signal from confocal sum projections, n = 3-4; mean ± SD; *p < 0.05, **p < 0.01, One-Way ANOVA with Dunnet’s multiple comparisons test, CTR vs TGFb, CTR vs TGFb + SB.

### Live imaging

Live phase contrast imaging was performed on ImageXpress XLS Micro High content imaging system (Molecular Devices). Live Calcein-AM imaging was performed by replacing culture medium with medium containing a mixture of 1:100 Calcein AM (Life Technologies #C3099) followed by an incubation of 20 min in a humidified incubator (37°C, 5% CO_2_). After incubation, the plate was transferred and imaged with an ImageXpress Micro XLS-C (Molecular Devices) confocal microscope with 10X magnification.

### Immunofluorescence staining

Fixation was performed by adding 50 μL of 3.7% formaldehyde (Sigma #252549) in each well except for the graft inlet (A2 in Figure 1B). An extra 50 μL were added to the left perfusion inlets. After a 15 min incubation, wells were washed thrice with PBS for 5 minutes and then left in PBS at 4°C or stained immediately. For staining, PBS was aspirated, and chips were blocked and permeabilized for 2 hours by adding PBS containing 1% Triton X-100 (Sigma #T8787) and 3% bovine serum albumin (BSA, Sigma #A2153) while placed on an OrganoFlow rocker. Primary antibodies were then prepared on ice in an antibody incubation buffer consisting of 0.3% Triton X-100 and 3% BSA in PBS. The following primary antibodies were used: mouse anti-CD31 (DAKO #M0823), mouse anti-MRP2 (Abcam #ab3373), goat FITC-conjugated anti-Albumin (Bethyl Laboratories #A80-229F), mouse conjugated AF488 anti-αSMA (R&D #IC1420G), goat anti-LYVE1 (R&D #AF2089-SP), rabbit anti VE-cadherin (Abcam #ab33168). Dilutions of primary antibodies were 1:100, except for CD31 which was diluted 1:50. 40μl of primary antibody suspension was added per well and incubated at RT on the OrganoFlow rocker overnight. Thereafter, the wells were washed 3x with PBS supplemented with 0.3% Triton X-100 (washing solution). Secondary antibodies were prepared next in the antibodies incubation buffer and added after the last washing step. The following secondary antibodies were used: donkey anti-rabbit Alexa 647 (Sigma #SAB4600177), donkey anti-mouse 647 (Life Technologies #A31571), donkey anti-goat 647 (Thermo Fisher Scientific #A21447), goat anti-rabbit 647 (Thermo #A-21244), goat anti-mouse 555 (Life Technologies #A21422), goat anti-rabbit 555 (Life Technologies #A21428), goat anti-mouse 488 (Life Technologies #A32723), goat anti-rabbit 488 (Life Technologies #A32731) and donkey anti-goat 488 (Thermo #A-11055). Secondary antibodies were always diluted 1:250. Two drops per mL of NucBlue (Invitrogen #R37605) and ActinRed (Life Technologies #R37112) were added to the secondary antibody mixture to stain for nuclei and actin expression respectively. In some experiments, BODIPY (Invitrogen #D3861) was added to visualize lipid droplets. 40μl of secondary antibody suspension was added to the wells. The plate was then incubated overnight at RT on the OrganoFlow rocker, after which it was washed 3x with the washing solution. Finally, PBS was added to all wells and the plate could be stored for a maximum of 4 weeks at 4°C in the dark. Single slice, maximum or sum intensity projections were acquired using the confocal Micro XLS-C High Content Imaging Systems (Molecular Devices) using 4X, 10X or 20X magnification.

### Image analysis

Image processing and analysis was executed in FIJI v.1.52. Relative fluorescence intensity units (RFU) of sum projections were obtained by measuring the average pixel intensity of the circular region corresponding to the graft chamber opening.

### Compound preparation and exposure

SB-431542 (Cayman Chemical Company #13031), Firsocostat ((MedChem Express #HY-16901) and Staurosporine (Sigma #S4400) were prepared as a 1000X stock solution in DMSO and stored at −20°C. For fatty acid exposures we used Palmitate Saturated Fatty Acid Complex (palmitate, Cayman Chemical Company #29558) and BSA-Oleate Monosaturated Fatty Acid Complex (oleate, Cayman Chemical Company #29557) which are both provided at a stock concentration of 5 mM. Recombinant Human TGF-β1 (TGF-β, Peprotech #100-021) was reconstituted to 20 μg/mL in 0.1% BSA and stored at −20°C. Compounds were dissolved in normal culture medium (EGM2 + Aprotinin and iHEP medium + Aprotinin) to their working concentration and added to the perfusion channels or graft chambers respectively. Checkerboard-patterned exposure layouts were dispensed using a Biomek i5 (Beckman Coulter). For the staurosporine exposure experiment, a mistake was made in which the cells were cultured in iHEP medium without glucose for the initial 24 hours of the experiment (Figure 5?).

### Enzyme-linked immunoassay (ELISA)

ELISAs for connective tissue growth factor (CTGF; R&D Systems #DY9190-050) and albumin (Sanbio #E88-129) were used to measure the secretion of the proteins in the medium of the graft chamber well following manufacturer’s instructions. Medium was collected prior to analysis and was stored in well plates at −20°C. For CTGF and albumin, samples were diluted 1:50 and 1:100 in the supplied diluent respectively. Absorbance was measured with the Spark® Cyto (Tecan).

### Robust Z’ factor calculation and optimization

To determine the robust Z’ factor, the following formula was applied:

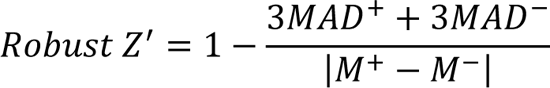

Where *MAD^+^* and *MAD^-^* refers to the median average deviation of the positive and negative controls multiplied by 1.483 and *M^+^* and *M^-^* refer to the medians of the positive and negative controls. We then followed a method that describes an iterative process where a new value for a datapoint is created by a weighted sum of the individual parameters (the individual assays, in this case normalized CTGF release, aSMA and actin signal) for that specific datapoint (individual chip). The weight for each parameter is then selected by maximizing the Z’ factor of the newly formed projection (32).

### Vasculature morphological assessment

Morphological descriptors were derived from confocal maximum intensity projections of liver cultures stained for the vasculature marker LYVE-1. For this, an automated workflow has been established using FIJI. First, images are pre-processed with a background subtraction algorithm. Then, the Labkit plugin (33) for FIJI was used to segment the vessels. This plugin enables for background and foreground annotations through training of a random-forrest pixel classification model to successfully segment the vessels as a binary image. Images are then postprocessed by removing structures outside the gel compartment and small artifacts. Explant area, vessel area, and vessel density are extracted from the binary image. The binary image is then skeletonized to extract the remaining morphological descriptors using FIJI’s built-in Analyze Skeleton function.

### Statistical analysis

Data were handled in Excel (Microsoft) and GraphPad 9. Analyses of variance (ANOVAs) were performed for multiple means comparisons with the built-in software provided by GraphPad. Significance was determined for p values < 0.05.

## Results

### Stellate cells and liver-derived endothelial cells self-organize in a vascular network

For this study, we used the OrganoPlate Graft, a microfluidic platform designed for the vascularization of spheroids and organoids (31). This platform consists of 64 microfluidic chips patterned underneath a standard 384 well-plate as illustrated in Figure 1A. Each chip consists of 6 fluidically-connected wells, with two lateral perfusion channels (From well A1 to B1 and from well A3 to B3) separated by Phaseguides (34) to a central channel (well A2 to B2) which is typically filled with a hydrogel (Figure 1B). All the channels are confined between two glass plates, with the exception of a circular opening in well B2, called the Graft Chamber. While the plate has been used to establish microvascular beds by induction of angiogenesis of lateral endothelial tubules (31,35), others have directly seeded endothelial cells (and supporting cells) within the hydrogel in the Graft Chamber observing a spontaneous self-assembly in microvascular structures (36). In this study, we followed a similar approach which is illustrated in Figure 1C. First, a mixture of liver-derived cells, fibrinogen and thrombin was prepared and directly seeded in the opening of the Graft Chamber. Following polymerization of the gel, human umbilical vein endothelial cells (HUVECs) were seeded in the lateral channels. The plate was then placed on a rocking platform that enables gravity-driven perfusion through the perfusion channels.

To assess the cell seeding density of primary bulk NPCs required to form self-organized microvascular structures, we seeded three different concentrations of liver bulk NPCs (5416, 8125 and 12188 cells / µL). On day 10, we observed the highest concentration of cells (12188 cells / µL) formed an extensive vascular network, connecting the two adjacent HUVEC tubules as visualized by VE-Cadherin in Figure 1D. Confocal microscopy revealed lumens within the vessels which are associated with alpha-smooth muscle actin (αSMA)-expressing cells (Figure 1E), indicating a role for hepatic stellate cells (HSCs) in liver-derived vascular bed formation in the OrganoPlate Graft. To evaluate this, we prepared three different cell mixtures containing 6500 primary liver-derived endothelial cells (LDECs) / µL with different amounts of primary HSCs (20:1, 4:1, 2:1). Indeed, at the lowest ratio (20:1), few and disconnected vessels were visible after 9 days of culture (Figure 1F, i) while with an increased number of HSCs (4:1 and 2:1), we observed a substantial increase in vascular structures (Figure 1F, ii and iii). Taken together, this demonstrates the importance of HSCs in vascular bed formation in the OrganoPlate Graft.

Next, we assessed whether HSCs within these cultures can be triggered with a fibrotic challenge. For this, we cultured LDECs or HSCs as monocultures or as co-culture in high density (LDEC:HSC 6500:3250 cells/µL) or low density (LDEC:HSC 3250:1625 cells/µL) for 7 days, followed by 24 hours of 10 ng/mL TGF-β with or without 10 µM SB-431542, a potent inhibitor of the TGF-β type I receptor (37). Confocal imaging revealed an increased expression of F-actin upon exposure to TGF-β which could be ameliorated by co-administration of SB-431542 (Figure 1G). This was evident for HSCs monoculture and co-culture, but not for LDECs monoculture. This indicates that HSCs within this system are capable of a fibrotic response, which was significant in both, HSC monoculture (data not shown) and in co-culture with LDECs (Figure 1H).

### Inclusion of hepatocytes results in liver microstructure formation

In the liver, hepatocytes account for approximately 80% of the total cell population and are responsible for the majority of the organ’s functions (38). In this study, we used iHEPs seeded alongside LDECs and HSCs at a density of 10’000 cells / uL hepatocytes, 3250 cells / µL LDECs and 1250 cells / µL HSCs (ratio 10:3:1). After 7 days, LDECs in triculture generated widespread vascular structures (Figure 2A) as indicated by the expression of LYVE-1 and CD31. An extensive population of albumin-positive cells in proximity of the vascular structures confirmed the presence of hepatocytes (Figure 2B). Higher magnification confocal imaging revealed a complex, but spatially organized cellular structure (Figure 2C and illustrated in Figure 2F). We observed lumenized microvasculature (black arrow) associated with stellate cells (white arrows) and hepatocytes (red arrows), as well as structures reminiscent of bile canaliculi between hepatocytes (blue arrow). Live-imaging of Calcein-AM confirmed close proximity of lumenized microvessels (red arrow) with hepatocytes (Figure 2D). Polarization of hepatocytes was confirmed by immunofluorescence staining of MRP-2 (Figure 2E).

**Figure 2:**
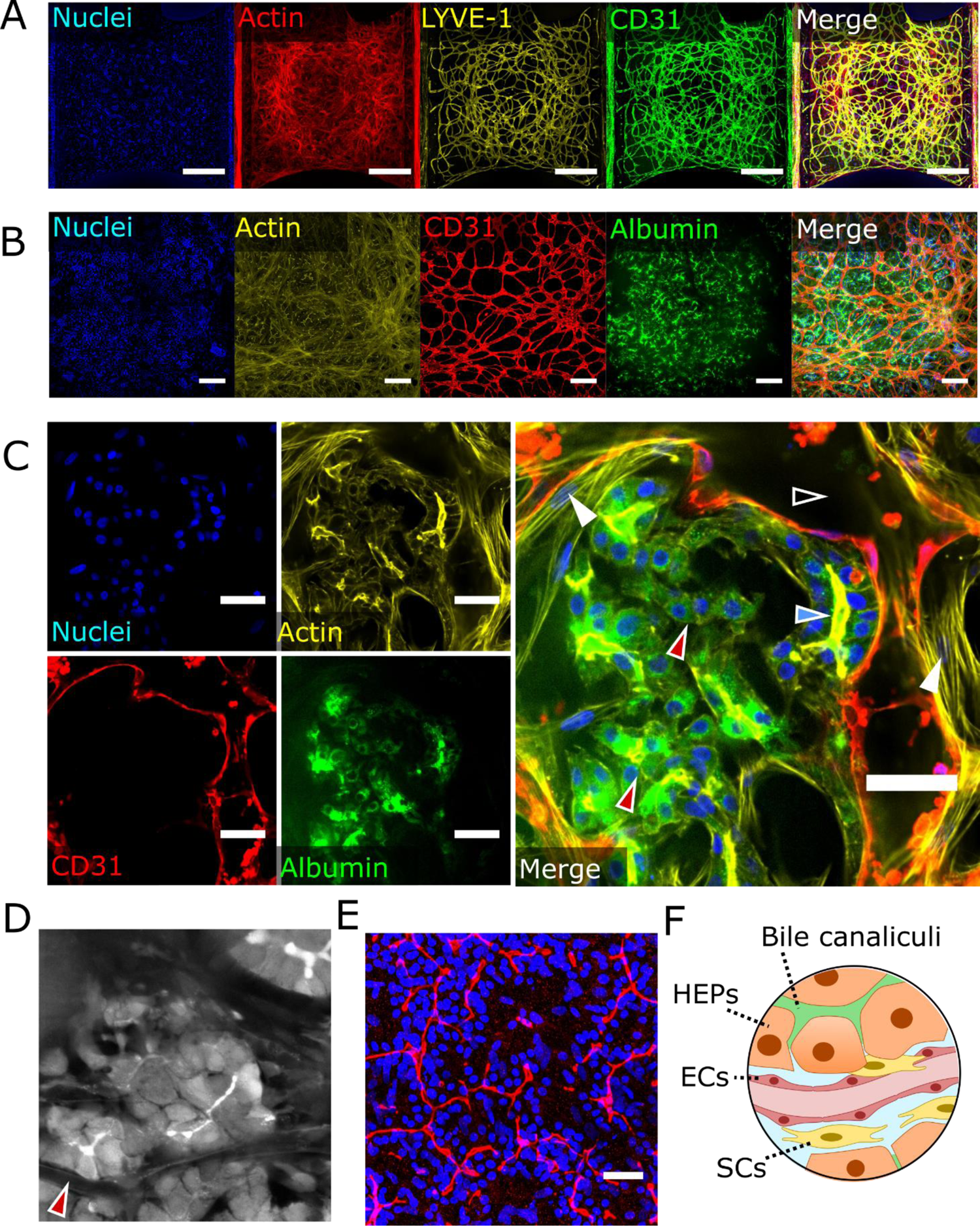
Integration of hepatocytes into liver microvascular system. **(A)** Confocal maximum intensity projections of a triculture stained for the vascular markers LYVE-1 and CD31, as well as F-actin and nuclei after 7 days of culture. Scalebar = 500 µm **(B)** Confocal maximum intensity projections of a triculture stained for the hepatocyte marker albumin, vascular marker CD31, F-actin and nuclei after 7 days of culture. Scalebar = 200 µm. **(C)** Confocal single slice stained for albumin, CD31, actin and nuclei. In the merged panel hollow vascular structures are visible (black arrow), stellate cells (white arrows) as well as hepatocytes (red arrows) and bile-canaliculi-like structures (blue arrow). Scalebar = 50 µm. **(D)** Live calcein-AM images showing hepatocytes associated with lumenized vessel (red arrow). **(E)** Confocal maximum projection image showing MRP-2 staining (red) and nuclei (blue). Scalebar = 50 µm. **(F)** Illustration of cell arrangement within the system.

### Steatosis can be induced in the comprehensive liver model

Excessive hepatic accumulation of lipids is one of the major leading causes of MASH (39). Supplementing hepatocyte culture medium with free-fatty acids (FFA), palmitic acid and oleic acid, is a common procedure to establish hepatic steatosis models (40). To demonstrate the feasibility of mimicking steatosis, a combination of palmitic acid (150 µM) and oleic acid (150 µM) was applied to the liver culture (iHEP, LDEC, HSC) for 12 days. Over the course of the incubation period, there was substantial darkening of the cells under phase contrast (Figure 3A). On day 12, BODIPY staining (Figure 3B) confirmed a significant accumulation of lipids within the culture (Figure 3C). The darkening of the culture as a result of lipid accumulation in the cells suggested that it would be possible develop a label-free assay to monitor lipid accumulation over time. To assess this, we performed an absorbance scan using a plate reader on an untreated chip and one supplemented for 12 days with 150 µM palmitic acid and 150 µM palmitic acid. Indeed we observed an increase in absorbance across the spectrum for the chip exposed to fatty acids (Supplementary Figure 1A). To demonstrate the feasibility of modeling steatosis prevention, cultures were exposed to FFAs with 2.5 or 10 uM Firsocostat, an acetyl-CoA carboxylase inhibitor which has been shown to reduce steatosis in a complex *in vitro* spheroid model as well as in clinical studies (12,41,42). We measured absorbance at 3,5,7 and 12 days after start of exposure at 900nm wavelength. On exposure day 5, there was a significant reduction in absorbance values of cultures treated with Firsocostat (10 µM) compared to controls. (Figure 3D). After 12 days of exposure, both concentrations showed significant reduction in absorbance measurement (Supplementary Figure 1B). Endpoint BODIPY staining confirmed significantly reduced intracellular lipid accumulation (Figure 3E) for both concentrations of Firsocostat (Figure 3F).

**Figure 3:**
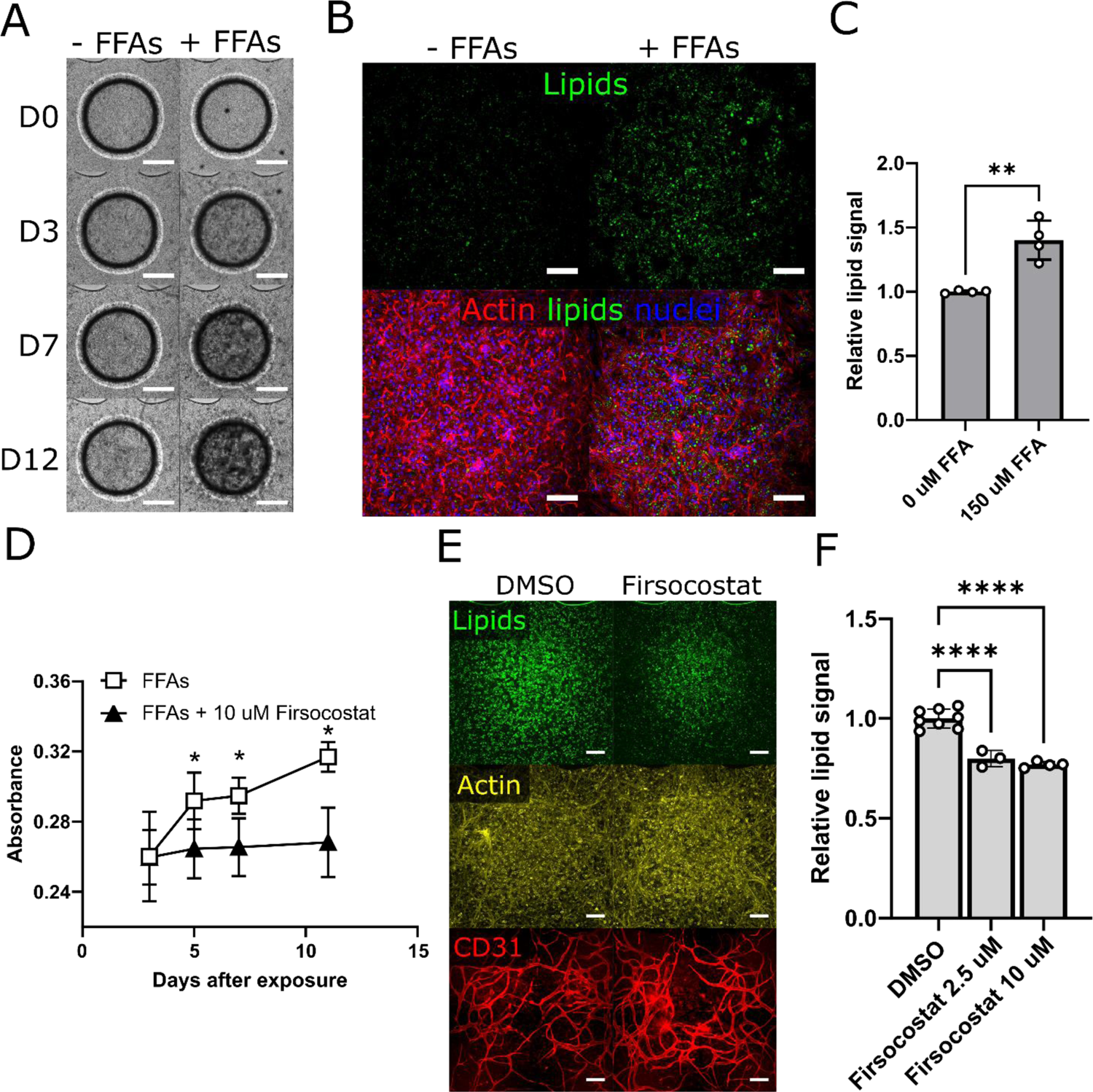
Development of a steatosis assay. **(A)** Phase contrast images over the course of 12 days exposure to FFAs. Scalebar = 500 µm **(B)** Confocal maximum intensity projections after 12 days of FFA exposure stained for lipids (green), F-actin (red) and nuclei (blue). Scalebar = 200 µm. **(C)** Confocal sum projections quantification of BODIPY signal, n = 3-4; mean ± SD; **p < 0.01; Unpaired T-Test, 0 uM FFA vs 150 uM. **(D)** Absorbance at 900 nm measured over the course of 12 days of chips exposed to FFAs with or without 10 µM Firsocostat, n = 3-8; mean ± SD; *p < 0.05; 2-Way ANOVA (Šídák’s multiple comparisons test), FFAs vs FFAs + 10 µM Firsocostat. **(E)** Confocal maximum intensity projections after 12 days of FFA exposure with or without Firsocostat stained for lipids (green), CD31 (red) and nuclei (blue). Scalebar = 200 µm. **(F)** Confocal sum projections quantification of BODIPY signal from chips exposed to FFAs with or without Firsocostat, n = 3-8; mean ± SD; ***p < 0.0001; 1-Way ANOVA, DMSO vs 2.5 or 10 µM Firsocostat.

### Development of an automated, multiplexed TGF-β-induced fibrosis assay

In the natural progression of the MASH, persistent fat accumulation leads to inflammation by activation of Kupffer cells (43). During this acute inflammation stage, activated Kupffer cells and other cells secrete TGF-β which is one of the most potent fibrogenic cytokines (44). TGF-β acts on quiescent HSCs which differentiate towards a myofibroblast phenotype, characterized by proliferation, ECM deposition and α-SMA(45). Connective tissue growth factor (CTGF), secreted by liver cells during fibrosis, is an important mediator and enhancer of TGF-β signaling and has been used as a biomarker for liver fibrosis both *in vivo* and *in vitro*(46,47).

Using CTGF secreted into the media as a primary readout, we optimized the assay by evaluating the optimal combination of TGF-β and SB431542. First, we assessed whether a lower TGF-β concentration could produce the same response as 10 ng/mL over the course of 72 hours. We observed significant increase in CTGF release in all tested concentrations (1, 5 and 10 ng/mL) with 10 ng/mL showing the highest response (Figure 4A). We confirmed, therefore, 10 ng/mL as the ideal concentration to produce the maximum response. Next, we investigated an optimal concentration of SB431542 that could be used to inhibit the response. For this, we cultured the chips for 72 hours with 10 ng/mL TGF-β with 0, 0.01, 0.1, 1 and 10 µM SB431542. We observed a reduction in CTGF release for all tested concentration, yet only 10 µM was capable to fully inhibit the fibrotic response (Figure 4B). We performed a vehicle tolerance test by the response of the treatment to increasing concentrations of DMSO. We found that the response did not significantly change for concentrations up to 1% DMSO, while 10% DMSO drastically influenced the assay (Figure 4C). We thus confirmed that the combination of 10 ng/mL TGF-β and 10 µM SB431542 provides the largest assay window as well as the absence of a vehicle-driven effect up to 1% DMSO. We repeated the experiment with an exposure of 10 ng/mL TGF-β and 10 µM SB431542 and we observed elevated F-actin and α-SMA protein expression that could be ameliorated by the addition of SB431542 (Figure 4D). This effect could be quantified using confocal sum projections for both F-actin (Figure 4E) and α-SMA (Figure 4F) as well as by confirming CTGF in the culture medium (Figure 4G).

**Figure 4:**
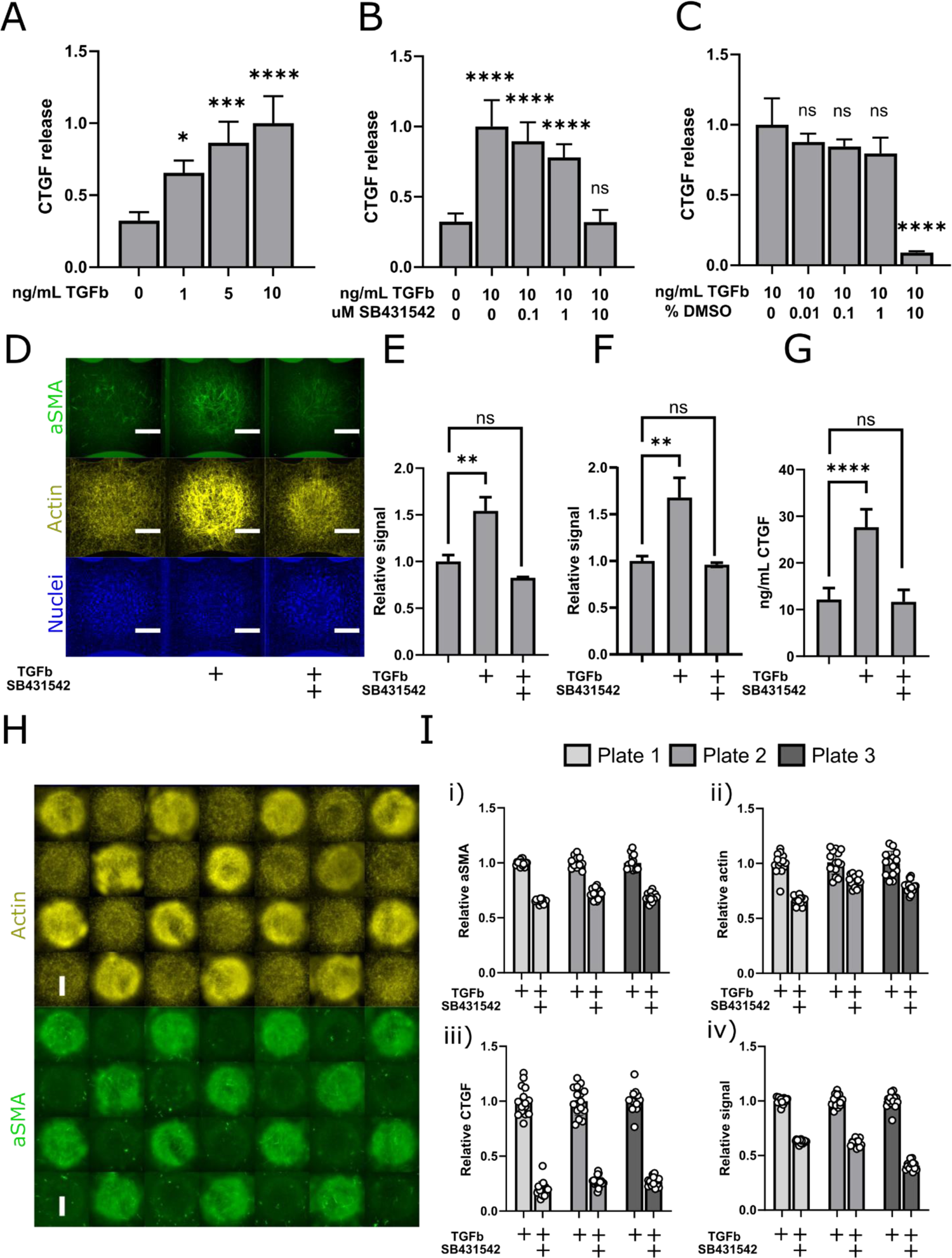
Development of TGF-β-induced screenable liver fibrosis assay. (**A**) Quantification of released CTGF in the culture medium after exposure. n = 8; mean ± SD; ****p < 0.0001, ns = not significant; 1-Way ANOVA, CTR vs TGFβ or TGFβ + SB. (**E**) CTGF release as a result of increasing concentration of TGF-β. n = 4-30; mean ± SD; *p < 0.05, ***p < 0.001, ****p < 0.0001; 1-Way ANOVA, Untreated vs 1 ng/mL, 5 ng/mL or 10 ng/mL TGFV. (**B**) CTGF release as a result of increasing concentration of SB431542. n = 4-30; mean ± SD; ****p < 0.0001; 1-Way ANOVA, Untreated vs 10 ng/mL TGFβ + 0, 0.1, 1 or 10 µM SB. (**C**) CTGF release as a result of increasing concentration of DMSO. n = 3-30; mean ± SD; ****p < 0.0001, ns = not significant; 1-Way ANOVA, 10 ng/mL TGFβ + 0, 0.01, 0.1, 1 or 10% DMSO. (**D**) Confocal maximum intensity projections showing aSMA, F-Actin, and nuclei, of triculture chips after 24 hours incubation with 10 ng/mL TGF-β with or without 10 µM SB431542. Scalebar = 500 µm. (**E**) Quantification of F-actin signal from confocal sum projections. n = 2-3; mean ± SD; **p < 0.01, ns = not significant; 1-Way ANOVA, CTR vs TGFβ or TGFβ + SB. (**F**) Quantification of aSMA signal from confocal sum projections. n = 2-3; mean ± SD; **p < 0.01, ns = not significant; 1-Way ANOVA, CTR vs TGFβ or TGFβ + SB. (**H**) Confocal sum projections of chips exposed to 10 ng/mL TGF-β with or without 10 µM SB431542 in a checkerboard pattern across the plate. Scalebar = 500 µm. (**I**) Reproducibility of fibrosis readouts across three different plates showing quantification of sum projection images from aSMA signal (i), F-actin (ii), released CTGF (iii) and combined readout (iv). For all comparisons, Dunnet’s multiple comparisons test was used.

We then automated the seeding and exposure procedures using a liquid handler. To assess the robustness of the automated fibrosis assay, on day 4, the complete model was exposed to TGF-β (10 ng/mL) with or without SB431542 (10 µM) for 72h in a checkerboard pattern across three different OrganoPlates, in two independent experiments. Staining for actin and α-SMA revealed strong differences in protein expression between the conditions (Figure 4H). Quantification of the staining together with CTGF released in the culture medium confirmed good reproducibility of the assays for all three readouts (Figure 4I i, ii, iii). To evaluate the Z’ factor of a high throughput, multiplexed screening campaign, we combined the three assays following a method that describes the integration of multiple readouts (32). When applied to the individual plates, this combined assay yielded a robust Z’ Factor for plate 1, 2 and 3 of 0.71, 0. 48 and 0.69 with an average 0.62 ± 0.13 (Figure 4I, iv) which is overall suitable for high throughput screening applications.

To identify the possibility of misidentifying toxicity as an inhibitory effect, we expanded the readouts to assess the distinctive signature of a known toxic compound, staurosporine, compared to SB431542 (summarized in Table 1). For this, we conducted a 10 ng/mL TGF-β exposure as described before but in presence of staurosporine (10 µM). Confocal microscopy revealed extensive loss of nuclear, aSMA, F-Actin and LYVE-1 signal after staurosporine exposure (Figure 5A). Both SB431542 and staurosporine caused significant decrease of CTGF release (Figure 5B), aSMA signal (Figure 5C) and F-actin signal (Figure 5D). Yet, staurosporine caused a 5-fold increase in LDH activity measured in the culture medium (Figure 5E) while SB431542 did not significantly affect this readout. Further, staurosporine significantly lowered and SB431542 increased the nuclei count (Figure 5F). To understand the impact of the compounds on hepatocyte viability, we quantified the amount of albumin released in the medium and observed a significant increase for SB431542 exposure while staurosporine exposure resulted in a decreasing trend, yet not statistically significant (Figure 5G). Finally, we employed an automated image processing method to extract shape descriptors from vascular networks. This revealed no significant effect of SB431542 yet strong detrimental effect of staurosporine on vessel density (Figure 5H), total number of junctions (Figure 5I) and vessel length (Figure 5J).

**Table 1:** Summary of readout outcomes in the TGF-β-induced fibrosis assay with inclusion of SB431542 (effective response) and staurosporine (toxic response) compared to DMSO.

| Readout | Significance | Effective response | Toxic response |
| --- | --- | --- | --- |
| CTGF release | Fibrosis | Decrease | Decrease |
| aSMA expression | Fibrosis | Decrease | Decrease |
| Actin expression | Fibrosis | Decrease | Decrease |
| Nuclei count | General viability | Increase | Decrease |
| LDH release | General viability | No change | Increase |
| Albumin release | Hepatocyte function | Increase | No change/decrease |
| Vessel density | Vascular morphology | No change | Decrease |
| Number of junctions | Vascular morphology | No change | Decrease |
| Vessel length | Vascular morphology | No change | Decrease |

**Figure 5:**
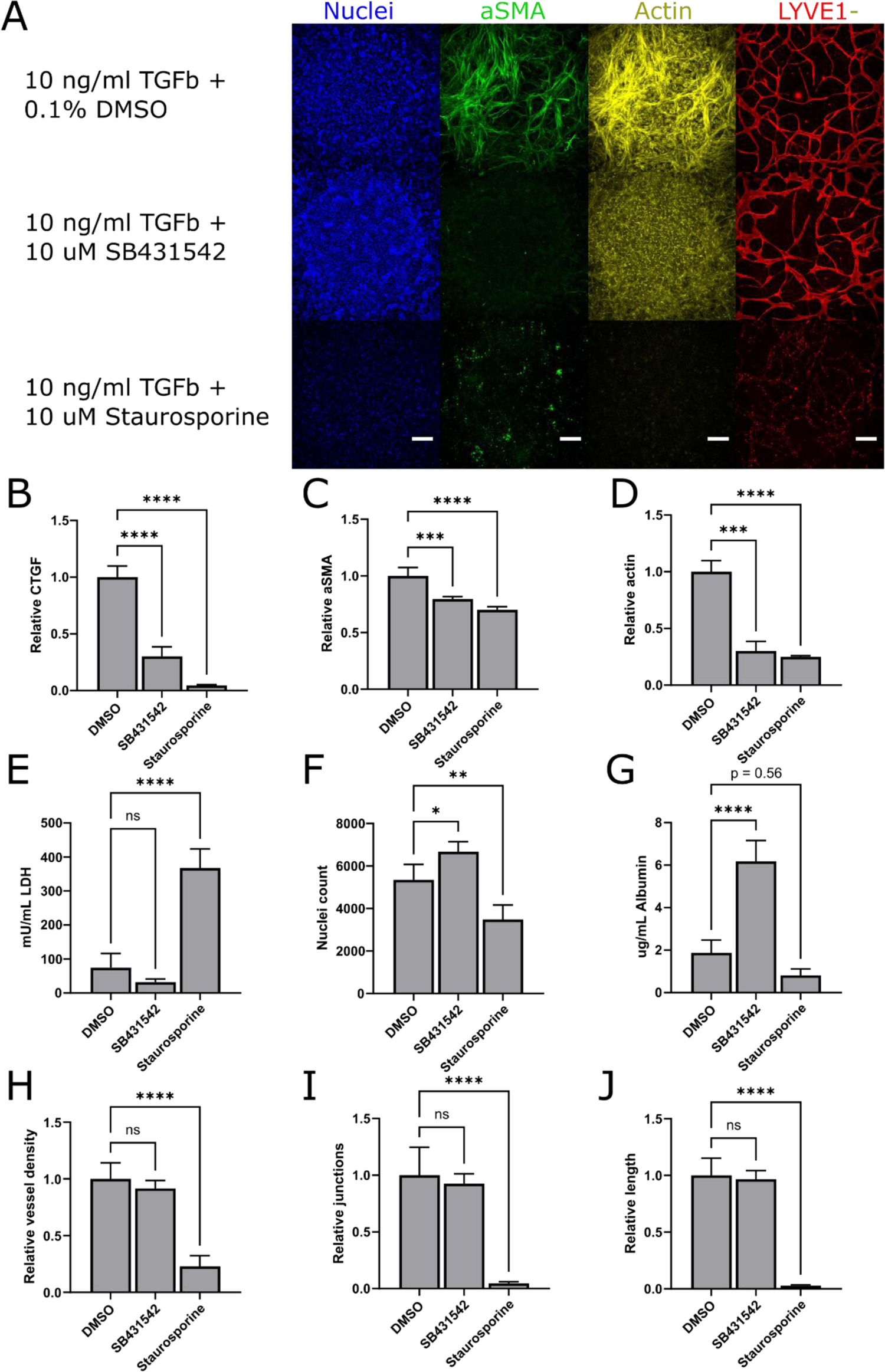
Toxicity signature on the fibrosis assay. (**A**) Confocal maximum intensity projections of triculture chips exposed for 72h to 10 ng/mL TGF-β in presence of 10 µM SB431542 or staurosporine (10 µM). Scalebar = 200 µm. (**B**) Quantification of released CTGF in the culture medium after exposure. (**C**) Quantification of aSMA signal from confocal sum projections. (**D**) Quantification of F-actin signal from confocal sum projections. (**E**) LDH activity assay on culture medium after exposure. (**F**) Nuclei count of the cultures after exposure. (**G**) Quantification of released Aabumin in the culture medium after exposure. (**H**) Relative vessel density after exposure. (**I**) Relative number of vascular junctions after exposure. (J) Relative vessel length after exposure. For all graphs: n = 4; mean ± SD; *p < 0.05, **p < 0.01, ***p < 0.001, ****p < 0.0001, ns = not significant; 1-Way ANOVA, Dunnet’s multiple comparisons test, DMSO vs SB, DMSO vs Staurosporine.

## Discussion

Representing human disease physiology in a non-clinical system with industrial applicability is the ultimate goal of MPS (48). Yet, these two aspects are often described as conflicting, increasing the fidelity of one aspect might consecutively decrease it in the other (49,50). Structural complexity, cellular composition, tissue stratification, fluid flow and extracellular environment are often achieved via sophisticated microfluidic designs with complex operating equipment which hinders ease of use and automation. Conversely, scalable systems often sacrifice resemblance of the organ’s natural state including the vascular component and are thus limited in their organotypic features and applications. Here, we describe a liver model that challenges this compromise with unprecedented organotypic features at the level of cellular organization, including an extensive vascular system, within a scalable and automated platform that allows for robust, multiplexed assays.

To achieve this, we leveraged the self-organization properties of cells embedded in extracellular matrices. This allows for a simplistic design of the culture chamber, such as the middle channel of the OrganoPlate Graft, where cells would spontaneously organize into structures including a vascular bed without a pre-conceived architecture. There are numerous reports of the self-assembly capacity of endothelial cells to form vascular networks in chip systems, including in (51–54). the OrganoPlate Graft in the context of a lung model (36). Yet, literature regarding liver-specific microvascular formation remains scarce. Here we demonstrate that bulk liver NPCs spontaneously form microvascular structures when embedded in a fibrin ECM. As it has been shown that the formation of endothelial networks can be modulated by a variety of stromal cell types (55–57), we hypothesized that, in a hepatic context, stellate cells would influence the emergence of vascular structures. Indeed, we found that hepatic stellate cells positively affected the number of microvessels suggesting high level of heterocellular interaction (Figure 1D and 1F).

While seeding a mixture of gel precursor, together with epithelial cells, endothelial cells and supporting cells has been reported as a viable strategy for establishing vascularized microtissues in microfluidic devices (51,58,59), this has only been partially explored in the hepatic context. Studies indicate that ECM encapsulation of pluripotent stem cell-differentiated hepatic cells improves function and cell maturation (60–62). Here we showed that seeding iHEPs alongside primary LDECs and HSCs leads to the formation of a complex, self-organized architecture that is reminiscent of the microscopic arrangement of liver cells found *in vivo*. We found that endothelial cells express the liver sinusoidal endothelial cell marker LYVE-1 and form extensive networks also in presence of hepatocytes, which undergo spontaneous re-arrangement (Figure 2A, C). This leads to a cellular disposition with hepatocytes closely interacting with both, stellate cells and endothelial cells, as well as forming apical structures reminiscent to bile canaliculi as indicated by MRP-2 expression (Figure 2E).

To demonstrate the utility of the vascularized liver model, we established robust assays for detecting processes involved in MASH including steatosis and fibrosis. The multifactorial nature of MASH and the intricate interplay between different cell types necessitates the establishment of complex, non-clinical models like the one reported here. Regarding steatosis, we established two assays capable of detecting increased intracellular lipid accumulation (Figure 3A-C), including a label-free detection method allowing for fast, real-time lipid accumulation monitoring, with minimal intervention and simplified quantification compared to imaging methods. Importantly, these readouts could be used to assess the pharmacological activity of a clinically-relevant compound. Firsocostat, an acetyl-CoA carboxylase, has been shown to reduce hepatic steatosis in spheroid models, animal models and in humans (12,41,42). We demonstrated that firsocostat significantly reduced the hepatic lipid accumulation in our model using both methods.

Following persistent hepatic fat accumulation, the next stage in the progression of MASH involves the onset of inflammation which promotes the activation of stellate cells responsible for the fibrogenic process. Among several factors, increased gut permeability is hypothesized to be involved in the development of MASH.

Microbial molecules reaching the liver activate Kupffer cells which release cytokines that activate stellate cells to promote fibrosis (43,63). A complete MASH model would, therefore, combine hepatic steatosis and Kupffer cell activation with subsequent fibrosis development as it has been shown in several *in vitro* studies (11,12,64). The model presented here does not include Kupffer cells and is therefore lacking this link. Instead, we triggered fibrosis by exogenous administration of TGF-β which is a key cytokine involved in the myofibroblast differentiation of stellate cells (45,64). TGF-β-induced fibrotic phenotype has been reported to be less relevant compared to free-fatty acid-induced fibrosis acids (64) suggesting that at this stage, our model may be limited in recapitulating the complete pathogenesis of MASH. Instead, we focused on establishing robust methodologies to assess the applicability of this platform for the measurement of fibrosis from a technical perspective. We automated the seeding procedures, optimized and multiplexed three fibrotic readouts: aSMA expression, F-Actin expression and CTGF release that were combined to maximize an assay window. With this, we demonstrated that despite its complexity in terms of cellular composition and architecture, this liver MPS model can be elevated with fully automated, multiplexed, reproducible assays with windows and throughput suitable for high-throughput screening (64 chips per plate with robust Z’>0.5).

Signal reduction in the fibrotic readouts also occur in the presence of toxic compounds, demonstrating the need to differentiate effective from toxic responses. Here, we multiplexed several readouts in order to distinguish between the inhibition of fibrosis and general or hepatocyte cytotoxicity. An effective response, as indicated by SB431542 showed reduction in fibrosis, no increase in LDH release, an increase in nuclei count and in albumin secretion and no changes in vascular morphology. On the other hand, general cell toxicant staurosporine showed, in addition to a reduction in fibrosis, a clear increase LDH release, a decrease in nuclei count, no increase in albumin secretion and drastic deterioration of vascular morphology.

A distinguishing aspect of this model is the morphological integrity and reproducibility of the vascular networks. We demonstrate here that this could potentially be used to assess liver vascular damage during drug development programs.

## Conclusion

In conclusion, we developed a fully vascularized liver MPS model comprising hepatocytes, hepatic stellate cells and liver-derived endothelial cells by leveraging the self-assembly capability of cells when embedded in a biological hydrogel. This resulted in a cellular architecture that shows unprecedented fidelity to the native organ. The model recapitulated key biological aspects of disease progression found in MASH. Concurrently, its robustness, as well as the underlying platform, likely permits its implementation for drug development as well as academic research. While the proposed model presents biological value, we envision the integration of the immune component as a crucial next step. The platform design, in combination with anastomosis between microvasculature and lateral vessels, allows for the integration of circulating cells which could potentially populate the liver tissue, elevating its applicability to include complex liver diseases, cancer and toxicity studies. In addition, utilizing the vasculature as a delivery route will likely gain relevance to predict efficacy of new drug modalities, such as biologics, viral vectors and cell therapy approaches.

## Acknowledgements

We would like to thank the Mimetas Biocore team including Arthur Stock, Manon Haarmans and Jeroen Heijmans for their technical support in automated procedures and cell banking services. We would also like to acknowledge the importance of Thomas Olivier and Aleksandra Olczyk for their role in supporting the development of the automated workflow to extract morphological features from stained microvessels. Flavio Bonanini received funding from the European Union’s Horizon 2020 research and innovation program under the Marie Skłodowska-Curie grant agreement No 812616. This project has received funding from the Innovative Medicines Initiative 2 Joint Undertaking (JU) under grant agreement No 945473 (IMI ARDAT). The JU receives support from the European Union’s Horizon 2020 research and innovation program and EFPIA.

## Data availability statement

The datasets used and/or analysed during the current study available from the corresponding author on reasonable request.

