## Supplementary Figure 1 for "A microvascularized *in vitro* liver model for disease modeling and drug discovery"

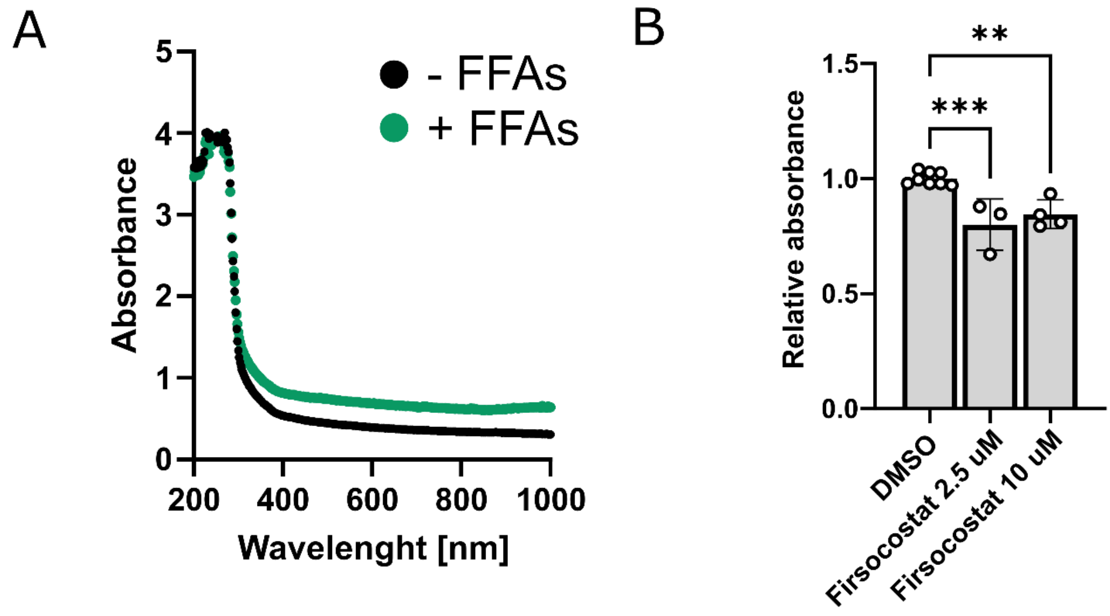


**Supplementary Figure 1: Label-free steatosis measurement (A)** Absorbance scan of two chips cultured for 12 days with (green line) or without (black line) FFAs. **(B)** Day 12 absorbance measurement at 900 nm of chips exposed to FFAs with or without Firsocostat. n = 3-8; mean ± SD; **$p < 0.01$, ***$p < 0.001$; Unpaired T-Test, 0 uM FFA vs 150 uM.
